# Boundary convection during velocity sedimentation in the Optima analytical ultracentrifuge

**DOI:** 10.1101/2021.03.08.434487

**Authors:** Steven A. Berkowitz, Thomas Laue

## Abstract

Analytical ultracentrifugation (AUC) provides the most widely applicable, precise and accurate means for characterizing solution hydrodynamic and thermodynamic properties. In recent times AUC has found broad application in the biopharmaceutical industry as a first-principle means for quantitatively characterizing biopharmaceuticals. Boundary sedimentation velocity AUC (SV-AUC) analysis is widely used to assess protein aggregation, fragmentation and conformational variants in the same solvents used during drug development and production. SV-AUC is especially useful for the analysis of drug substance, drug product and dosing solution, where other techniques may exhibit solvent matrix issues or concentration limitations. Recently, the only manufacturer of the analytical ultracentrifuge, released its newest (third generation) analytical ultracentrifuge, the Optima, in early 2017 to replace its aging 2^nd^ generation XL series ultracentrifuges. However, SV-AUC data from four Optima units used in conducting characterization work on adeno-associated virus (AAV) has shown evidence of sample convection. Further investigation reveals that this problem arises from the temperature control system design, which is prone to producing destabilizing temperature induced density gradients that can lead to density inversions. The observed convection impacts both the qualitative and quantitative data generated by the Optima. The problem is intermittent and variable in severity within a given Optima unit and between Optima units. This convection appears to be mainly associated with low rotor speeds and dilute samples in dilute solvents, such as AAV samples in formulation buffers containing relatively low concentrations of salts, sugars, etc. Under these conditions it is found that a sufficiently robust stabilizing density gradient is not always present during sedimentation, making the sample susceptible to convection by localized density inversions. Because SV-AUC is used as an analytical tool in making critical decisions in the development and quality control of biotherapeutics, it is imperative to alert users about this potential problem. In general special attention to data quality needs to be made by those researchers working with very large biopharmaceutical particles (e.g. gene therapy products that involve viral vectors or nanoparticles), where the conditions leading to convection are most likely to occur. It is important to note that the XL series analytical ultracentrifuges do not suffer from this problem, indicating that this problem is unique to the Optima. Attributes that reveal the presence of this problem and strategies for its elimination or minimization are provided.

## Introduction

Despite being nearly 100 years old, analytical ultracentrifugation (AUC) continues its role as a useful and important analytical tool in both basic and applied sciences [1–11]. Its longevity and modern capabilities, recently summarized by Peter Schuck and coworkers in a comprehensive three volume series [12–14], is a result of being the most accurate and precise technique for characterizing a solution’s thermodynamic and hydrodynamic properties. The availability of absorbance, refractive and fluorescence detection allows AUC’s application to a broad range of samples and solvents. Size distribution analysis by boundary sedimentation velocity AUC (SV-AUC) offers access to a wide range of sample characterization information that includes the following: structural heterogeneity (e.g., fragmentation, aggregation and chemical composition) [13], concentration [15], interactions [14], changes in conformation [16] and sample consistency [10] with few solvent matrix limitations, minimum sample manipulation and easy method development [7]. Importantly, AUC readily offers significant and unique possibilities for characterizing the new classes of very large, complex molecular biopharmaceutical gene therapy delivery systems that are now being actively developed, such as viral vectors [17,18], liposomes and nanoparticles [19–25].

AUC is a first-principle technique where data interpretation does not require any comparison with standards. This means that, to within a percent or two, measurements made on a sample on one machine by one user at one time should be reproducible made “effectively” on the same sample at any other time by another user on a different machine. Any deviations between the two sets of measurements (that is beyond instrumental uncertainties) must reflect differences in the samples. Accuracy is required of every component of the machine to achieve a high certainty in such differences [26–31].

In an analytical ultracentrifuge there are several critical elements requiring high precision and accuracy: 1) sample cell geometry and alignment in the rotor, 2) machining of the rotor cell hole positions and sizes for radial position determinations, 3) temperature measurement and control, 4) optical measurement systems, 5) rotor speed control and 6) timing. Except for the actual cell alignment in the rotor, AUC users depend on the instrument manufacturer’s ability to make sure these critical elements (and their constituent components) deliver the required accuracy and precision. Temperature accuracy and control is of critical importance for the proper interpretation of SV-AUC data due to its impact on solvent density and viscosity (at 20° C, a 1 degree temperature change results in a 4% sedimentation coefficient change) and to avoid adverse density gradient situations that can lead to convection [27–29,32–39].

During sedimentation a stabilizing density gradient (i.e. increasing solution density with radius) builds up due to solvent component (e.g. salts, buffer and sugars) redistribution. If this stabilizing gradient is weak, as occurs with dilute solvents at low rotor speeds, the sample (especially dilute samples) becomes extremely sensitive to density inversion that can lead to convection. A key driver of density inversions in AUC arises from temperature gradient conditions in the sample that give rise to a condition where a cooler, denser, volume element (dV) is centripetal to a warmer, less-dense volume element (see Figure 1). Such a situation occurs when positive radial temperature gradients exist in the sample sector of an AUC cell. In the presence of a gravitational field the higher density region will be driven into the lower density solution resulting in the bulk exchange of material between the adjacent regions, as indicated in Figure 1.

**Figure 1.**
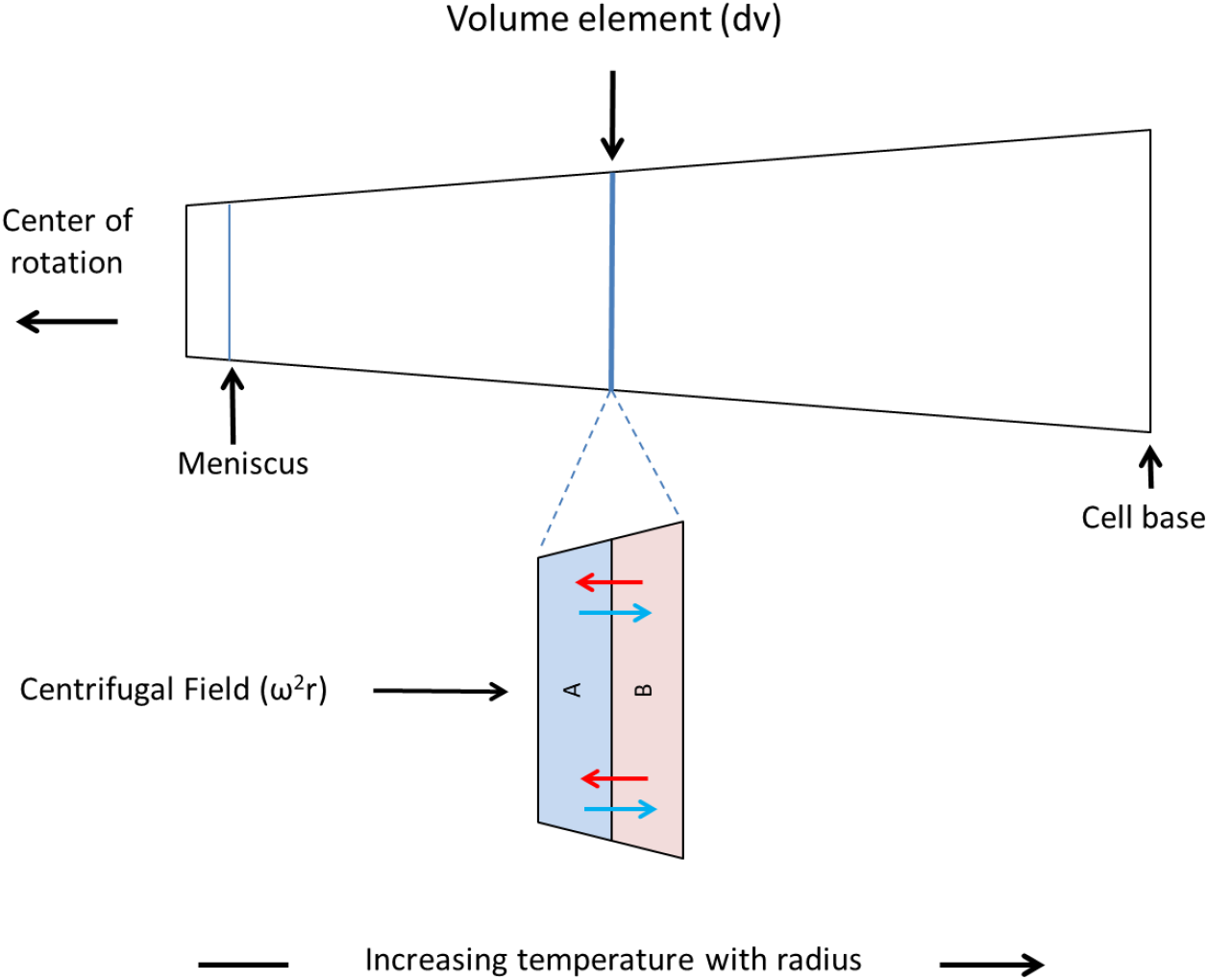
Illustration showing how the presence of a positive temperature gradient with increasing radius, within the sample sector of an AUC cell, can cause convection. Since the density of aqueous solutions decreases with increasing temperature, if the temperature of a sample (or some region of the sample) is a positive increasing function of radius a negative increasing density gradient with radius will be present. On looking at a volume element, dV, where this positive temperature gradient exists, a higher density region (A), located at shorter radius, above a lower density region (B), located at longer radius, will exist. Under these conditions in the presence of a gravitational field solution instability will exist leading to density inversion, where the solutions in A and B will exchange (as indicated by the blue and red arrows), leading to convective mixing in order to achieve a stable configuration in the gravitational field. However, if a sufficient positive increasing (stabilizing) density gradient with radius can be formed by solution solutes (macromolecules, salts, sugars, etc.) during centrifugation to override the unstable negative temperature induced density gradient, convective mixing will be avoided. Nevertheless, if such stabilizing density gradient cannot be achieved or cannot be achieved in time relative to the migration of the macromolecule solutes being studied, convection can occur.

The result of a density inversion is turbulent flow with concomitant mixing of the two adjacent solutions - convection. Historically, proper temperature control has been a challenging part of AUC’s development, where even the presences of a very small positive temperature gradient can lead to density inversion and sample convection due to the amplification effect of the gravitational field [32–39].

Recently (in early 2017) the premier manufacturer of analytical ultracentrifuges launched its third-generation analytical ultracentrifuge called the Optima, which were preceded by the Model E (1949 – 1972), and XLA/I (1990 – 2017) series analytical ultracentrifuges. In upgrading any instrument it is anticipated that the newest generation instrument will performs as well as, if not better than, its predecessors. Today this is even more important than ever given the application of AUC in the development of biopharmaceuticals, where it is employed in critical decision making. Unfortunately, early Optima units released into field exhibited a technical problem, see Berkowitz and Philo [10], associated with convective mixing during SV-AUC. This problem was uncovered when one of the authors (S.A. Berkowitz), working with one of the first users of the Optima, lead to information being reported back to the manufacturer indicating that while normal sedimentation patterns were obtained for a monoclonal antibody (mAb) when it was run at 40K rpm, no sedimentation was observed at lower rotor speeds (10-15K rpm), even after many hours of centrifugation. On receiving this information the manufacturer determined that the problem was due to a lack of temperature uniformity resulting in thermally-induced convection during centrifugation. To correct this problem the manufacturer introduced a thermal shroud within the rotor chamber along with some software changes to minimize thermal gradients in the rotor. With these modifications, it appeared that the Optima thermal convection problem was resolved.

However, even after these remedies were introduced into the Optima, subsequent work by S.B. Berkowitz in collaboration with several other Optima users has revealed intermittent artifacts in data scans from a number of Optima units during SV-AUC experiments involving adeno-associated virus (AAV). These artifacts are typically characterized by a spontaneous and unusual sharpening of the leading edge of the fastest moving boundary after migrating approximately quarter to almost half of the way through the sample sector. Data presented in this paper supports the conclusion that these artifacts are thermally-driven form of convection that still occurs in Optima units, arising most likely from the physical location of the Peltier heating/cooling cells at the rotor’s edge. By considering the key criteria for achieving convection-free SV-AUC (see Supplementary Material), recommendations are made for preventing this convection. These recommendations should be considered short-term measures for dealing with the situation until a permanent instrumental fix to the Optima can be achieved.

## Material and Methods

### Adeno-associated virus

All AAV samples used in SV-AUC experiments were produced and purified at the site associated with the specific Optima centrifuges outlined in the section heading “Analytical ultracentrifugation” below.

### Analytical ultracentrifugation

Optima AAV SV-AUC experiments conducted in this work were carried out on 4 different Optima units. They are designated as Optima_1, Optima_2 and Optima_3 & Optima_4. In addition, at one Optima site AAV experiments were also conducted on a XL-I. All AAV SV-AUC runs were carried out at a rotor speed of 18K rpm, using an An-50 Ti 8-hole rotor at 20°C and at AAV concentrations of about 1 OD or less at 230nm unless otherwise indicated. All AAV samples used in this work were placed in formulation buffers that contain physiological levels of salt, buffer, etc. totaling in the range of about 0.1-0.2 M at a pH near neutrality. All data analysis was carried out via SEDFIT, version 15.01b [40] using continuous c(s) distribution model where it is assumed that all species migrate independent of each other under ideal conditions.

In processing Optima interference data using SEDFIT standard corrections for radial- and time-independent (RI and TI) noise were made. However, in processing Optima absorbance data, the standard use of only time-independent noise correction, as done routinely in the case of the XL-A/XL-I, was found to be enhanced by including the RI-independent noise correction. This difference is believed to be due to the difference in how data acquisition is conducted between these two analytical centrifuges. In the case of the XL-A/XL-I absorbance detector, data acquisition for each sample first collects the reference intensity reading at radial position r, followed by the corresponding sample intensity reading at a slightly different radius (when using the ‘continuous scan’ mode). Meanwhile, the Optima acquires all intensity readings from the sample sector first, followed by the acquisition of all corresponding intensity readings from the solvent reference sector for the same range of radial positions. The resulting enhanced time delay between sample and reference solvent intensity readings in the Optima may allow for slight offsets to occur that enable the use of radial-independent noise correction to be useful to include in the case of processing absorbance data when using the Optima.

### Simulating AAV, mAb and LMW formulation buffer SV-AUC experiments

Simulations of SV-AUC experiments were conducted using SEDFIT using stated molecular weights, sedimentation coefficients, rotor speeds and data intervals of time between data scans. In all cases radial data spacing of 0.001 cm, and signal/noise (S/N) of 1 part 10^4^ at a starting arbitrary concentration unit of 1 was used. All simulations include the rotor acceleration phase.

## Results

### Unique abrupt data scan sharpening artifact in characterizing AAV material via SV-AUC using the Optima and its likely association with convection

SV-AUC experiments on AAV material using Optima_2 has revealed the intermittent appearance of boundary disturbances, see Figure 2A. These disturbances appear as a limited, but continuous block of data scans whose presence, absence or intensity varies both on a run to run basis, for a given Optima and between Optima units. Most often the disturbances, which are believed to be artifacts, are associated with the fastest moving boundary, appearing suddenly after the boundary has migrated about a quarter to half of the way down the sample sector. When observed, disturbances are frequently seen in all the cells in the given run. In some cases these anomalous disturbances can be very irregular in appearance; however, in most cases the disturbance can be summarized as a very abrupt boundary sharpening effect (associated with the leading edge of the boundary) that then rapidly broaden and disappears over subsequent data scan, see Figure 2A.

**Figure 2.**
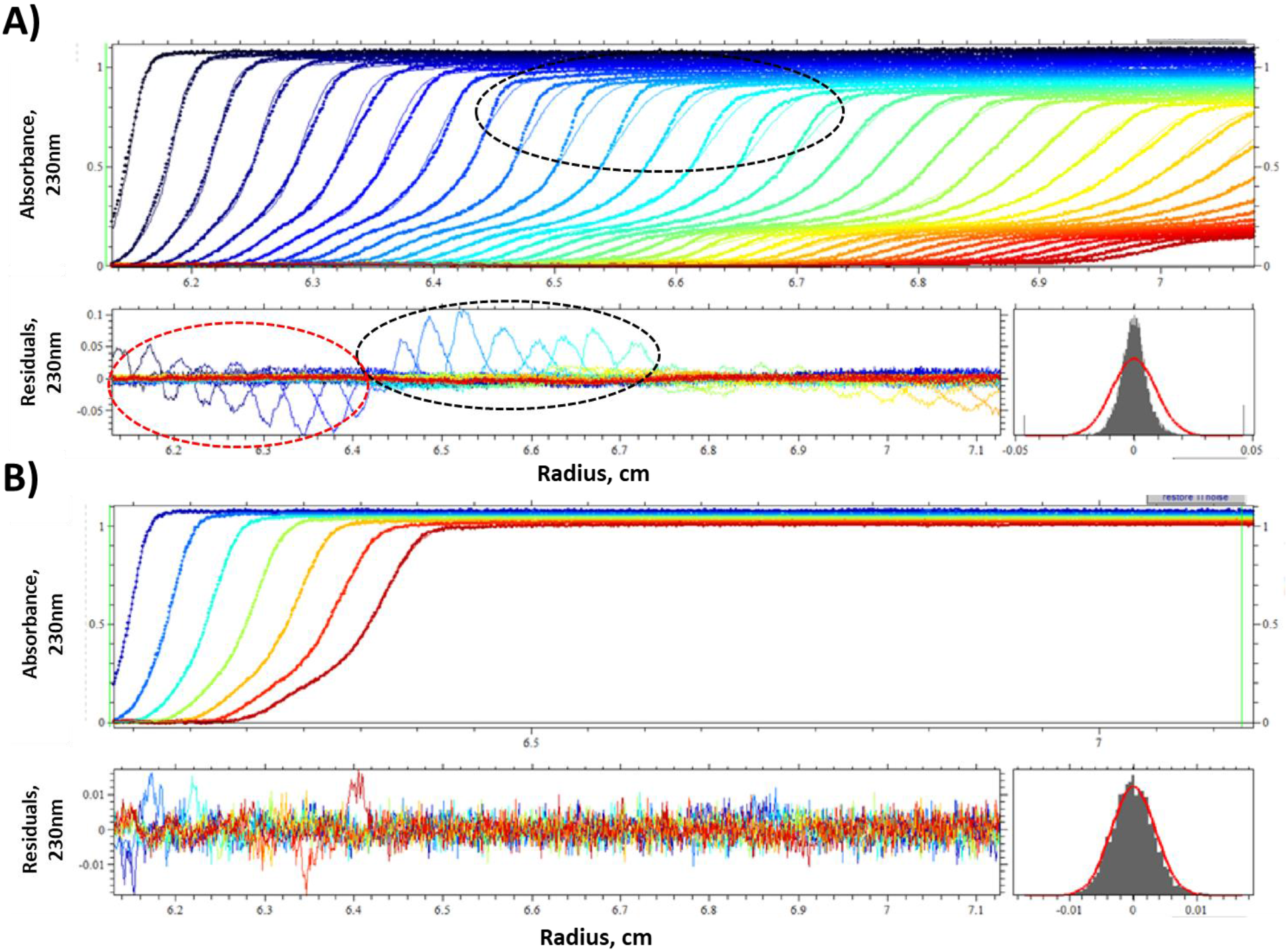
AAV SV-AUC data obtained from an Optima_2 run at 20K rpm and 20°C showing the presence of data scan sharpening artifact and evidence for its association with convection. **A) Top** plot is the overlay of the processed absorbance data scans 1-35, which encompasses the entire run**, bottom left** plot is the resulting residual plot showing the quality of the computed c(s) model fit to the experimental data shown in the top plot and the **bottom right** plot is the distribution of the experimental residuals (shown in the residual plot) relative to their anticipated normal distribution based on the data’s RMSD (outlined in red). Resulting statistical parameters from the c(s) analysis involving data scans 1-35 is the following: RMSD = 0.0093, Z-test = 110 and H = 19.9%. Areas highlighted by a black dashed oval in the overlay of data scans and residual plot indicate the region and block of data scans that visually show the presence of the data scan sharpening artifact (due to convection), while the red dashed oval in the residual plot indicates the region and block of data scans (1-7) that “*falsely*” show a poor fit of the experimental data to the c(s) model due to the *later* appearing convective disturbance in data scans. **B) To**p plot is the overlay of just the processed absorbance data scans 1-7 from the same data scan set shown in “A”, **bottom left** plot is the resulting residual plot showing excellent improved quality of fit of the c(s) model to the experimental data scans relative to that seen above in “A” (note the expansion in the Y-axis of this plot relative to the same plot shown in “A”), **bottom right** plot is the distribution of the experimental residuals (shown in the residual plot) relative to their anticipated normal distribution based on the data’s RMSD (outlined in red), which now shows an excellent fit to the normal distribution. Resulting statistical parameters from this c(s) analysis also significantly improved: RMSD = 0.0034, Z-test = 15 and H = 0.9%.

Further analysis of the AAV SV-AUC data shown in Figure 2A using just the earliest block of data scans acquired before the first visual signs of data scan sharpening are detected (what we refer to as “partial data scan analysis”) provides some additional data supporting the conclusion that the observed data scan sharpening is in fact a result of convectional disturbances developed sometime *later* after the AAV SV-AUC run is initiated. This is demonstrated by comparing c(s) analysis using all 35 data scans, Figure 2A vs c(s) analysis using just the first 7 data scans, Figure 2B.

In Figure 2A a poor fit to c(s) modelling is seen when all 35 data scans, which covers the entire sedimentation process of the sample. However, it turns out that the poor fit seen in the first 7 data scans is not due to any actual alterations or disturbances in these early data scans, e.g., due to convection. These early data scans are actually perfectly good (undisturbed) data scans that are miss fitted during c(s) analysis, when the entire set of 35 data scans are included, due to alterations in the *later* data scans that were impacted by convection (which abruptly appears *after* the first 7 data scans were acquired). This conclusion is revealed by separately analyzing the first 7 data scans (which were generated before the first sign of data scan sharpening is seen), independent of the remaining (altered) data scans, see Figure 2B. Using this partial data scan analysis approach, significantly improved c(s) modelling is achieved as indicated by improvements in the data scan overlay plot, residual plot, and the good agreement in the residual distribution plot to the expected residual distribution plot for a normal distribution of residuals having the same root mean square deviation, RMSD, [34]. In addition, the set of statistical parameters associated with the quality of the c(s) analysis model fit to the experimental data are also greatly improved, which includes the following: 1) RMSD, 2) H value (a comparison index of the experimental distribution of residuals, generated from the analysis of the best model fit to experimental data, yielding a given RMSD, relative to expected distribution of residual based on a normal distribution having the same RMSD [41]) and 3) runs test Z value (the number of standard deviations by which the runs of positive and negative residuals differ from the expectation of runs for a situation where residuals are normally distributed [41,42]) presented in the figure legend to Figure 2.

In cases where the Optima used had both absorbance and refractometric detectors (Optima_1), these disturbances are observed in the data collected by both detectors from the same sample, at the same radii and at the same time, as seen in Figure 3A and B. Similar results with another Optima, where the residual deviations were a factor of 2-3 times greater has provided conclusive data to support the conclusion that both detectors observe the same scan disturbances (data unavailable to be shown). Given the orthogonal nature of these two detectors it’s highly unlikely that the observed same data scan disturbances could be explained as a detector artifact. Rather the cause of these disturbances must lay in events occurring in the samples, as appose to the detectors. Similar anomalies have been observed in two other Optima units (Optima_3 and Optima_4).

**Figure 3.**
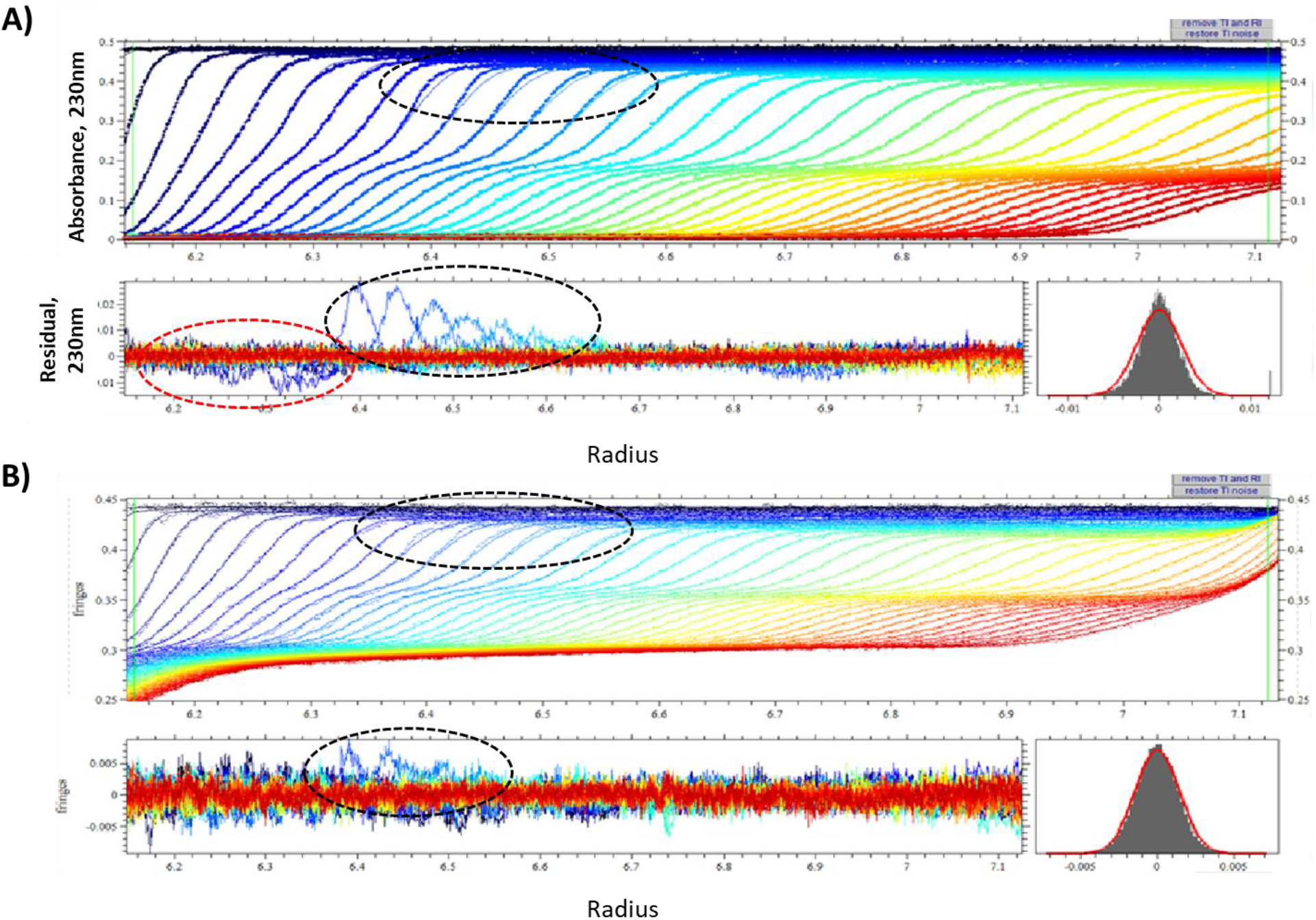
AAV SV-AUC data obtained from an Optima_1 run at 15K rpm and 20°C showing the presence of data scan sharpening artifact when data is collected on the same sample using both **A)** absorbance data at 230nm and **B)** corresponding interference data. In each case the **top** plot corresponds to the overlay of processed experimental data scans, while the **bottom left** plot is the resulting residual plot showing the quality of the computed c(s) model fit to the experimental data scans (shown in the top plot) and the **bottom right** plot is the distribution of the experimental residuals (shown in the residual plot) relative to their anticipated normal distribution based on the data’s RMSD (outlined in red). Area highlighted by a black dashed oval in the overlay of data scans and residual plots for both the absorbance and interference detector indicate the radial region in the block of data scans that visually show data distortion (artifact) due to convection. Area in the absorbance residual plot highlighted by the red dash oval correspond to the data region that “*falsely*” show a poor fit of the experimental data to the c(s) model due to the later appearing convective disturbance in data scans, as was similarly indicated in data shown in Figure 2.

While concentration-dependent hydrodynamic non-ideality may result in boundary sharpening [12,34,36,43], its effect occurs at concentrations orders of magnitude greater than that employed in AAV SV-AUC experiments illustrated here in Figures 2 and 3 (involving Optima_2 and Optima_1, respectively) and seen in work involving Optima_3 and Optima_4. Furthermore, the Optima aberrations appear abruptly some amount of time *after* the rotor has reached speed and then disappear over a few subsequent scans (Figures 2A and 3). Consequently, concentration-dependent hydrodynamic non-ideality cannot explain the Optima AAV data scan sharpening.

Importantly, it should be noted, no such artifacts are observed for AAV samples run under the same experimental conditions using an AUC instrument with different temperature control design (XL-I). Even samples that showed the artifact when run on the Optima did not show the artifact when run on the XL-I. Consequently, the problem is isolated to the unique design characteristics of the Optima. What is also important to note about the artifact is its adventitious nature. Slight variations in thermal component properties and/or operation may cause the observed variation in frequency and intensity of the artifact’s appearance, both in a particular Optima unit and between Optima units. Thus, in some cases the presence of the artifact can be quite subtle.

### SV-AUC Simulations of AAV and its formulation buffer solutes: Assessing qualitatively the ability of the AAV formulation buffer density gradient to achieve convection-free AAV SV-AUC

To achieve convection-free sedimentation requires the radial density gradient (dρ/dr) in the sample sector (for all appropriate radii, r, and time, t, values during SV-AUC) to be ≥ 0 (preferably > 0) (as outlined in the Supplemental Material). Since the concentration of AAV samples analyzed when using the Optima are very dilute, the low molecular weight (LMW) co-solutes (e.g., salts, buffer, sugar, etc.) that constitute the AAV’s formulation buffer, play the dominate role in providing stabilizing dρ/dr values to achieve convection-free sedimentation. However, given the relatively low rotor speed at which AAV SV-AUC experiments are conducted, combined with the rapid sedimentation properties of AAV material relative to the dynamics in developing the formulation buffer’s density gradient (via the redistribution of the LMW components in the formulation buffer during AAV SV-AUC) may not provide adequate stabilizing dρ/dr for all appropriate r and t values during AAV SV-AUC. If true, situations could exist during SV-AUC where migrating AAV material could be exposed to adverse destabilizing factors that could allow convection to occur.

Since the radial density gradient, dρ/dr, formed by any material run in an ultracentrifuge is directly proportional to the change in its concentration with the change in radius, dc/dr, (see discussion in Supplemental Material, *Key criteria for stable convection-free SV-AUC*), simulating the sedimentation process by generating the theoretical data scans (concentration as a function of radius) at the same series of time points for AAV and its formulation buffer and then comparing these two data scans at each time point can provide a quick qualitative insight into assessing the vulnerability of the AAV material to potential convection.

Using the SV-AUC stimulation capabilities in SEDFIT with approximate molecular weight (MW) and sedimentation coefficients values for the two key AAV species found in AAV samples, empty (of about 3.9 MDa and 62 S) and full (of about 5.2 MDa and 95 S) virus particles, and formulation buffer (using an approximate weight-average MW of about 100 Da and an experimental measured weight-average sedimentation coefficient of about 0.2 S,), theoretical ideal sedimentation data scans for these materials for the same time intervals and radial positions, can be computed (for the same sedimentation conditions, 18K rpm and 20°C, commonly used to conduct AAV SV-AUC experiments on the Optima). The resulting data scan comparisons from these calculations (for AAV and its formulation buffer components) for the same 6 time points (spaced 10 min. apart) are shown in Figure 4A-F.

**Figure 4.**
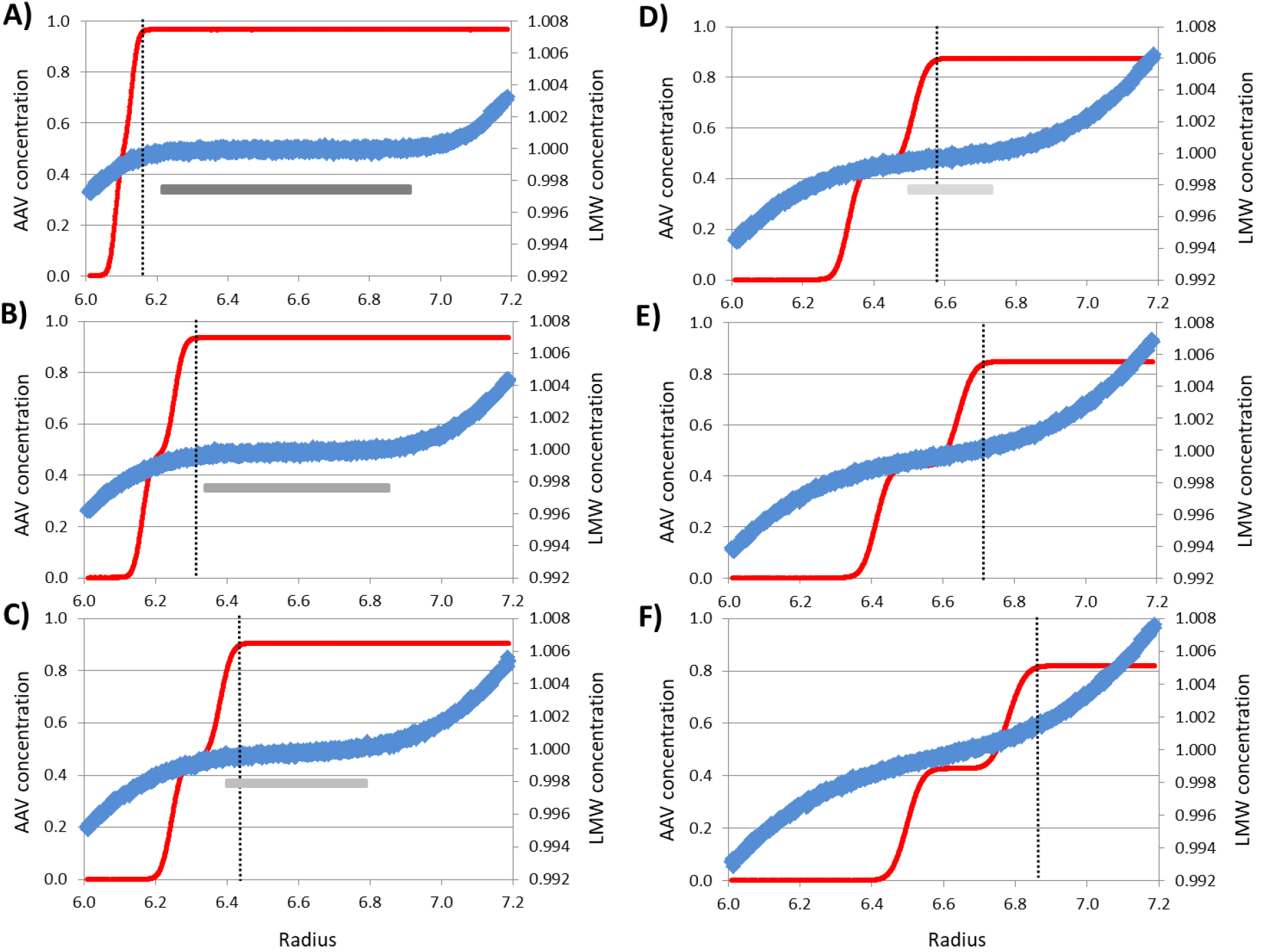
Simulated SV-AUC data for a hypothetical AAV sample consisting of a 50:50 mixture of empty and full virus particles, having a MW of about 3.9 MDa and a sedimentation coefficient of about 62 S, and a MW of about 5.3 MDa and a sedimentation coefficient of about 95 S respectively (in red), overlaid with the corresponding simulated SV-AUC data for a typical AAV formulation buffer system having a weight-average MW of 100 Da and sedimentation coefficient of 0.2 S (in blue), at 18K rpm and 20°C. Data scans for both the AAV mixture and formulation buffer were calculated for the times intervals of 10 min yielding data scans for the following time points (from the start of the run): **A)** 10 min, **B)** 20min, **C)** 30 min, **D)** 40 min, **E)** 50 min and **F)** 60 min for the same radial positions. Black vertical dotted line corresponds to the position of the leading boundary edge of the full AAV material. In both cases the initial starting concentration was an arbitrary concentration of 1. The length of the horizontal grey bars included in these plots indicates the extent of the radial region where the formulation buffer density gradient is the shallowest (where the potential for convection is most significant); while the lighter the grey the less prominent is the instability (or weakness) in this density gradient.

These simulations show that during the early phase of sedimentation the positive protective formulation buffer density gradient is only significant at the extremes of the AAV sample in the AUC cell’s sample sector (near the meniscus and cell base). Over time the formulation buffer density gradient propagates its way (via sedimentation and diffusion) throughout the entire AAV solution in the AUC cell to provide a final equilibrium density gradient. However, before this stabilizing density gradient can be fully established there is a dynamic changing radial time window, indicated by the grey scaled horizontal bars of different lengths in Figure 4 (where the darker the grey the weaker is the radial density gradient) that shortens with time during AAV SV-AUC, where this density gradient protection from the formulation buffer is very weak. For the fastest migrating AAV material (complete full virus particle) the migration of this material into any of these radial regions exposes this material to limited protection from adverse disturbing effects, such as negative density gradients, that can easily induce convection (e.g., from the presence of positive temperature density gradients with increasing radius). This situation appears to be most severe during approximately a 10 minute radial time window between the 30 and 40 min data scans acquired after the start of the AAV SV-AUC run, see Figures 4C and D. Qualitatively this corresponds to the observed intermittent time-window of artifacts in the experimental data.

For comparison purposes, similar simulations were carried out using the same formulation buffer, containing a monoclonal antibody, mAb, sample solution containing a 50:50 mixture of monomer (having a MW of about 150 kDa and a sedimentation coefficient of about 6.5 S) and dimer (having a MW of about 300 kDa and a sedimentation coefficient of about 9.8 S) run at 40K rpm at a temperature of 20°C. Results for these calculations are shown in Figure 5.

**Figure 5.**
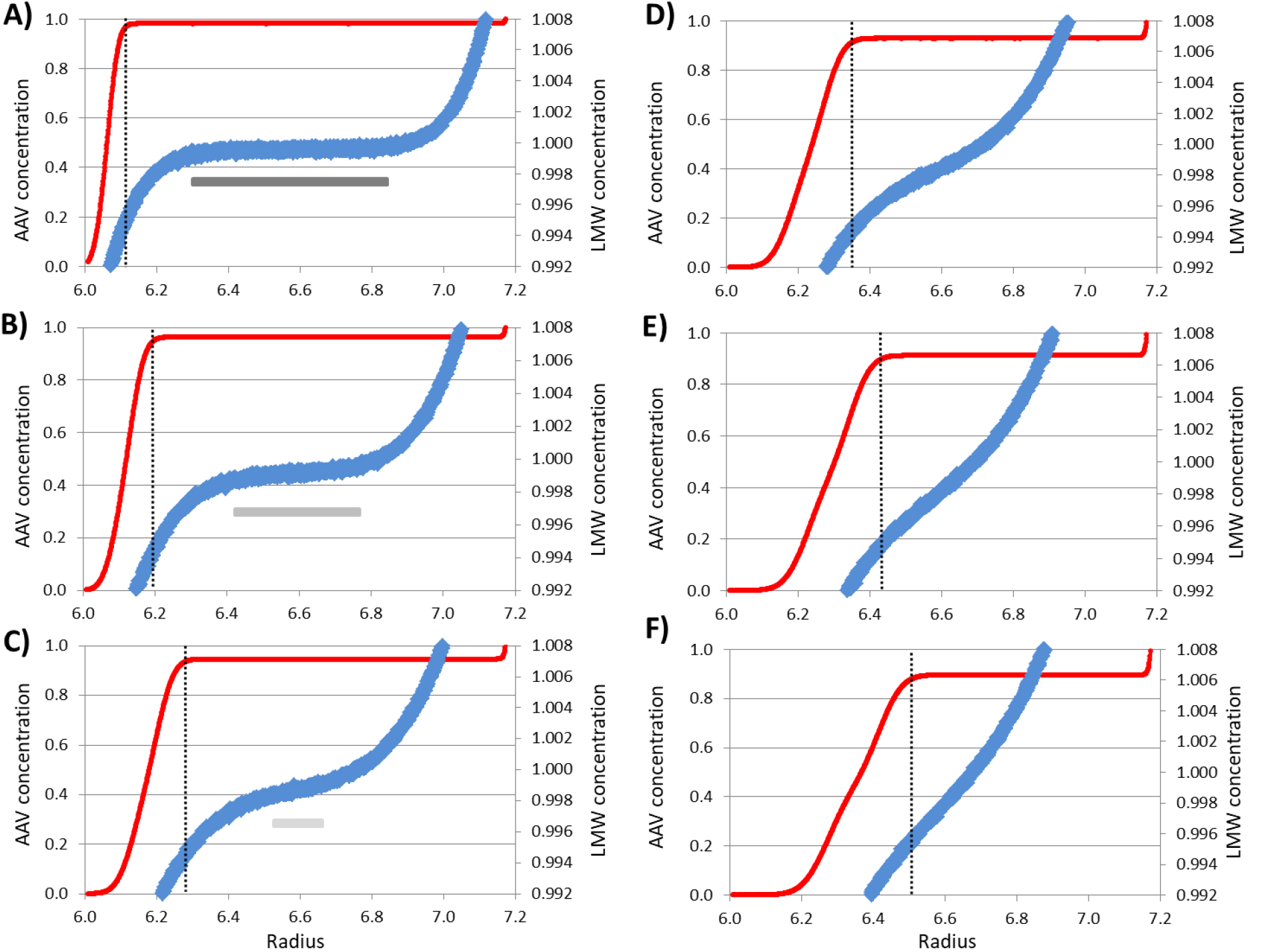
Simulated SV-AUC data for a hypothetical mAb sample consisting of 50:50 mixture of a mAb monomer and dimer, having a MW of about 150 kDa and a sedimentation coefficient of about 6.5 S and a MW of about 300 kDa and a sedimentation coefficient of about 9.8 S respectively (in red), overlaid with the corresponding simulated SV-AUC data for a typical mAb formulation buffer (which should be very similar to an AAV formulation buffer) having a weight-average MW of 100 Da and sedimentation coefficient of 0.2S, (in blue), at 40K rpm and 20°C. Data scans for mAb mixture and formulation buffer were calculated for the time intervals of 10 min, yielding data scans for the following time points (from the start of the run): **A)** 10 min, **B)** 20 min, **C)** 30 min, **D)** 40 min, **E)** 50 min and **F)** 60 min for the same radial positions. Black vertical dotted line corresponds to the position of the leading boundary edge of the mAb dimer material. In both cases the initial starting concentration was an arbitrary concentration of 1. The length of the horizontal grey bars in plots A-C indicates the extent of the radial region where density gradient is the shallowest (where the potential for convection is most significant); while the lighter the grey the less prominent is the instability (or weakness) in this density gradient. The absence of grey bar in plots D-F indicates the lack of any significance instability being present.

Data clearly shows that during the entire mAb SV-AUC run all of the mAb material migrates in the presence of a strong density gradient in compromise to the AAV SV-AUC simulated data (note in particular, that the Y-axis for the formulation buffer in both Figures 4 and 5 are scaled the same).

### Detrimental effect of a positive radial temperature gradient during SV-AUC and a comparison of the physical layout of the heating/cooling elements of the temperature control system employed by the three generations of analytical ultracentrifuges built by the same manufacturer

A key source for failing to achieve dρ/dr ≥ 0 (preferably > 0) during SV-AUC is the generation of opposing negative density gradients resulting from the presence of a positive radial temperature gradient (dT/dr), as discussed in the Supplement Material, under the heading *Detrimental effects of a positive radial temperature gradient during AUC*. As a result, if the radial formulation buffer density gradient, dρ_B_/dr, cannot override these adverse negative radial temperature induced density gradients, dρ_T_/dr, as indicated by eq. 6 in the Supplemental Material, convection-free SV-AUC experiments will not be achieved. Consequently, over the years a considerable amount of work has been given to designing appropriate temperature control systems for analytical ultracentrifuges that minimize the generation of such positive radial temperature gradients during AUC.

In the developing of the Model E, proper temperature control was achieved by cooling the outer rotor chamber wall with refrigerator cooled coils while heat was applied from the bottom of the rotor via a nichrome heating wire located at a radius between the radial position of the rotor cell holes and the rotor’s center of rotation, see Fig. 6A.

To maintain the AUC cell to a fixed desired running temperature and to maintaining a stable dρ/dr environment for AUC experiments a slight *negative* dT/dr gradient with increasing radius was maintained by cooling the outer rotor chamber to a slightly lower temperature than the desired running temperature to help avoid generating any destabilizing *positive* dT/dr gradients.

In developing the XL-A/XL-I both cooling and heating were provided from the rotor chamber bottom floor via Peltier cooling/heating cells, as indicated in Figure. 6B. In this case, the perpendicular nature of the direction of temperature gradients relative to the direction of sedimentation should minimize opportunities for generating adverse (positive) dT/dr gradients, and thus (negative destabilizing) density gradients that could cause convection.

In developing the Optima, however, the Peltier cooling/heating cells that control temperature are located radial along the outer rotor chamber wall (as cooling/heating towers), see Figure 6C. Given their radial position, any thermal output is guaranteed to generate positive (destabilizing) dT/dr gradients, especially during heating, that can propagate their way radially towards the center of rotation into the spinning sample solution during SV-AUC, see Figures 7.

**Figure 6.**
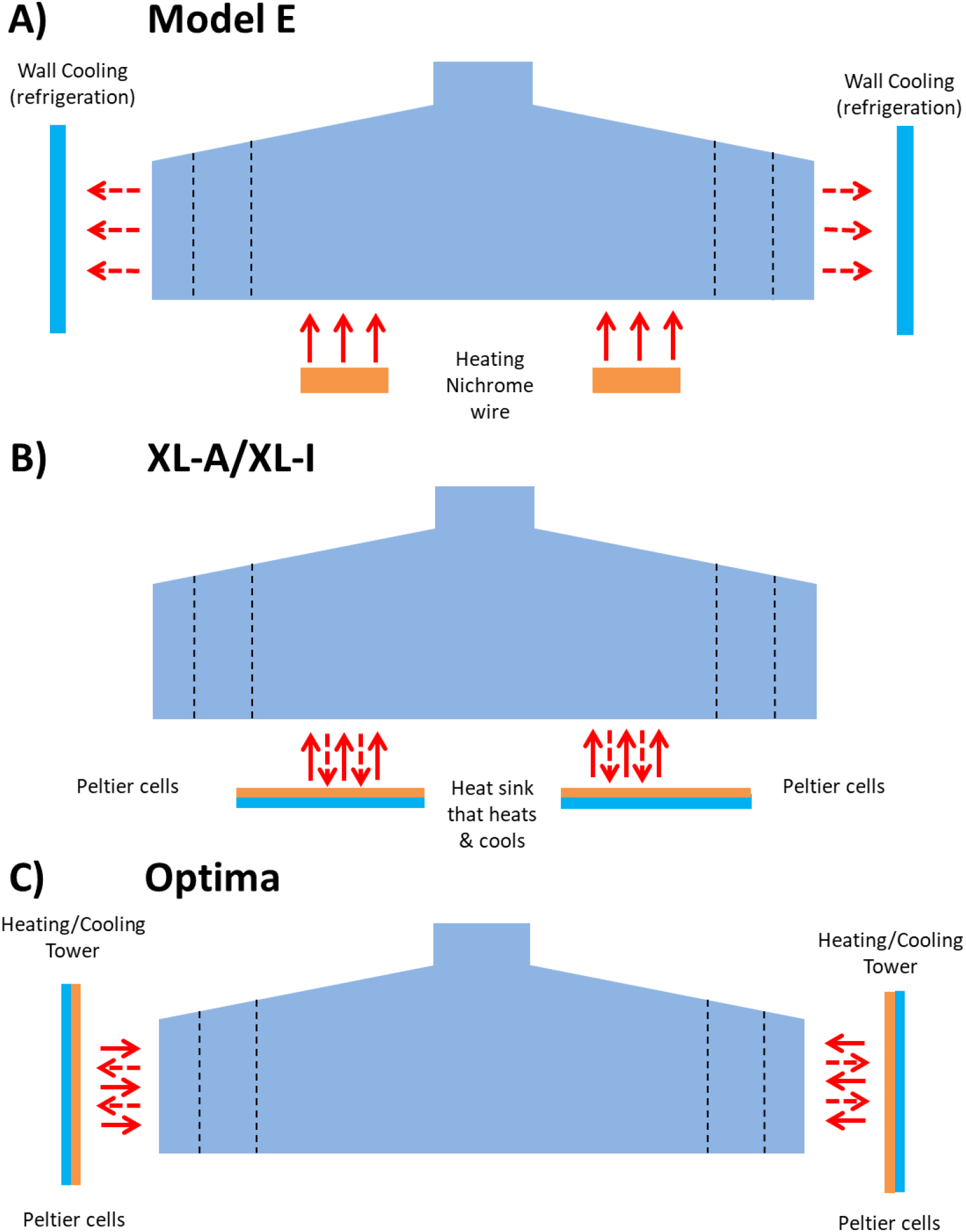
A coarse pictorial side view of the general layout of the temperature control system from past and present analytical ultracentrifuges made by the same manufacturer. **A)** Model E analytical ultracentrifuge, bottom area shows the approximate location of the nichrome wire heating element (orange horizontal bars at the bottom of rotor) and the refrigerated cooling coils located along the outside wall of the rotor chamber, indicated by the blue vertical bars, **B)** the XL-A/XL-I analytical ultracentrifuge, bottom area shows the rough location of the Peltier cooling/heating cells (indicated by the orange/blue horizontal bars at the bottom of rotor located between the center of rotation and the radial position of the rotor holes), and **C)** the Optima analytical ultracentrifuge with the cooling/heating Peltier cells located along the outside wall of the rotor chamber (indicated by the orange/blue vertical bars). In all cases the solid red arrows indicate the directional flow of heat during heating while the dashed red arrows indicate the directional flow of heat during cooling.

**Figure 7.**
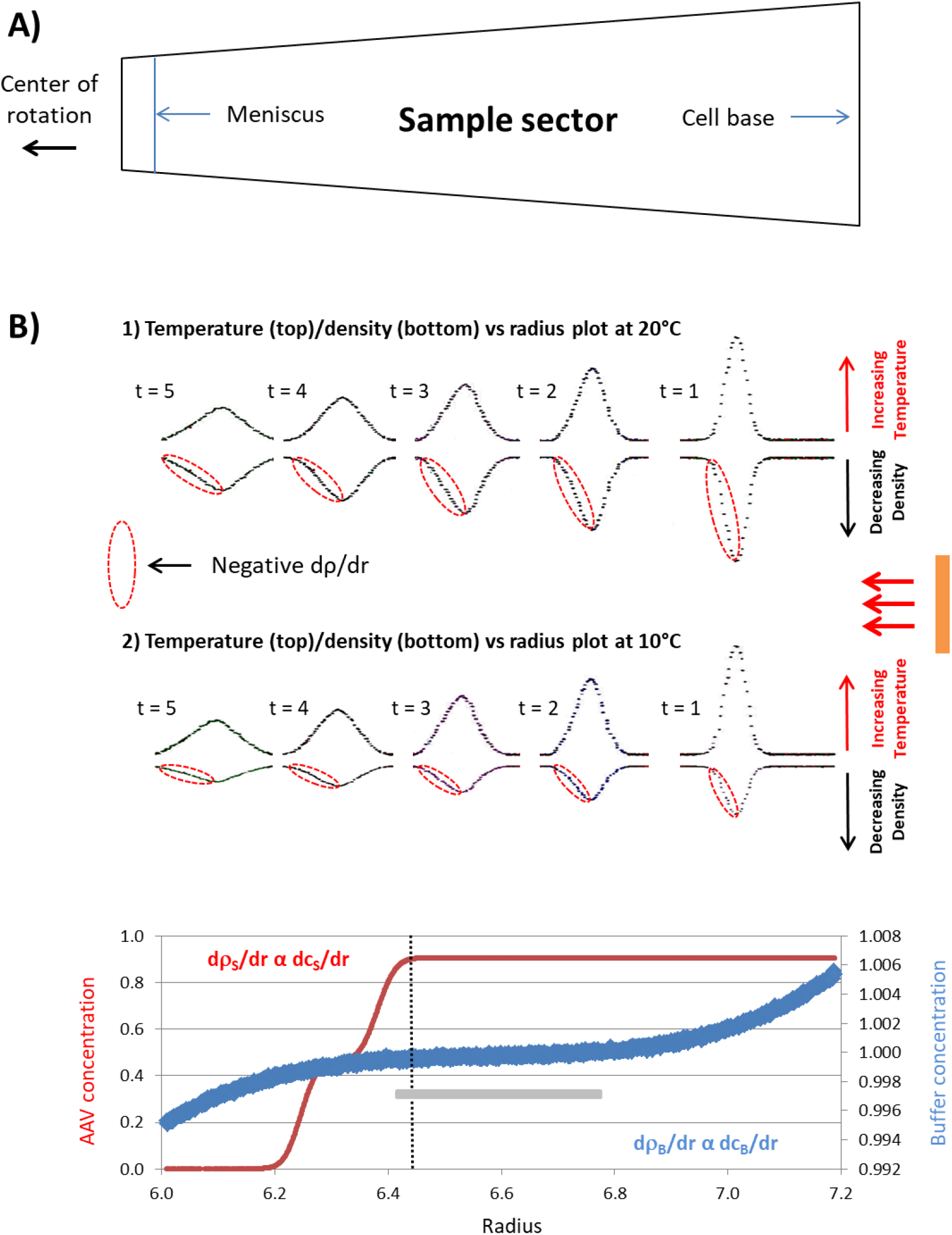
A summary slide showing how the present Optima’s temperature control system can induce convection during SV-AUC, via the introduction of positive radial temperature gradients (dT/dr) that in turn induced negative density gradients (dρ_T_/dr), and how conducting SV-AUC experiments at 10°C vs 20°C can eliminate or minimize the extent of convection (as discussed in the Supplemental Material, under section *Detrimental effects of a positive radial temperature gradient during AUC*). **A)** Is a pictorial view of the sample sector looking down at the AUC cell from the top of the rotor. **B)** For each temperature (parts 1 and 2) the **Top** plot portraits the progressive movement of the correspond identical hypothetical, intermittent output (illustrated as a heat pulse of some duration) from the Optima’s Peltier cells (located along the outer rotor chamber walls, indicated by the vertical orange bar) as it migrates radially towards the rotor’s center of rotation (indicated by the red arrows), as a function of time, t, through the sample sector of the AUC cell while the **Bottom** plot corresponds to the resulting impact on the sample solution’s density as a function of radius for the same time points. All plots for 10°C and 20°C were roughly radial scaled to the sample sector dimensions shown in part “A”. Note the red dashed ovals highlight the dynamic changing *negative* radial (destabilizing) density gradient (dρ_T_/dr) resulting from the corresponding dynamic changing *positive* radial temperature gradients (dT/dr). The results show that for the *same* temperature change, running at 10°C vs 20°C reduces the magnitude in the change in solution density significantly, which reduces the opportunity for convection. **C)** Same simulated overlaid calculations of the resulting sedimentation data scan of a 50:50 sample mix of empty and full AAV particles (S), in red, and it corresponding formulation buffer (B), in blue, after 30 min of sedimentation at 18K rpm at 20°C, as shown in Figure 4C. Note, this plot was roughly radial scaled to the sample sector dimensions shown in part “A”. Results show that at this time point into the AAV SV-AUC run the leading edge of the full AAV particle is exposed to the negative destabilizing density gradient propagating into the sample sector from the heat output of the Optima’s Peltier cells (located along the outer wall of the rotor chamber, which can over power the weak countering positive radial density gradient from the formulation buffer, dρ_B_/dr, (indicated in the radial region highlighted by the horizontal grey colored bar, which is the only significant source of density gradient that can override the negative temperature density gradient). Hence depending on the arrival and magnitude of negative temperature density gradient, dρ_T_/dr, relative to formulation buffer density gradient convection will occur if the negative dρ_T_/dr is greater than the positive dρ_B_/dr.

This situation can expose samples within their AUC cells to unstable negative dρ/dr conditions during SV-AUC, which is most critical when these negative density gradient encounter sample macromolecular solutes in radial regions where the formulation buffer density gradient is very weak, as shown in Figures 4C-D and 7C. Under such conditions if dρ/dr ≥ 0 (for all appropriate r and t values) is not met convection will result (through density inversion). A temperature control system, as outlined for the Optima, clearly places *significant excessive dependence* on having an adequate positive dρ_B_/dr gradient to override the presence of any negative dρ_T_/dr gradients to maintain stable convection-free SV-AUC conditions (dρ/dr ≥ 0).

### The effects of temperature on AAV data scan sharpening when conducting SV-AUC using the Optima

To acquire additional information to further support temperature as the key source of micro-convection seen in Optima during AAV SV-AUC, AAV SV-AUC experiments were conducted at 10°C and 30°C. At the lower temperature two things occur that should help suppress convection. The first, as the temperature of a solution is reduced from 20°C to 10°C the rate of change in dρ_T_/dT, decreases (in Supplemental Material see Figure 1S and discussion concerning eq. 7). In fact at 10°C dρ_T_/dT is nearly half that at 20°C (and is effectively zero near 0 - 4°C). This significantly reduces the change in density that one will see at 10°C relative to 20°C for the *same* small change in temperature, as illustrated in Figure 7B. Hence, conducting SV-AUC experiments at 10°C reduces the magnitude of strain put on the formulation buffer to provide an adequate positive dρ_B_/dr gradient to overcome the same positive temperature gradient. And secondly, at 10°C the viscosity of an aqueous solution is significantly greater than at 20°C. This latter effect should dampen any convectional turbulence that may arise.

Meanwhile, in the case of SV-AUC at 30°C, dρ_T_/dT is about 50% larger than dρ_T_/dT at 20°C (see Figure 1S in Supplemental Material). In this situation the impact of a temperature change on an aqueous solution’s density is significantly greater relative to the same temperature change at 20°C. As a result, at 30°C a greater demand is placed on dρ_B_/dr to provide an adequate density gradient to overcome a larger negative dρ_T_/dr than that would have formed for the same change in temperature at 20°C. Finally, at 30°C the viscosity of aqueous solutions is less than that at 20°C, reducing the positive benefits of viscosity to dampen the effects of convection.

Results from these temperature experiments, using Optima_4, which was found to nearly always show signs of data scan sharpening when conducting AAV SV-AUC runs at 20°C, revealed the presence of the typical signatures of the presence of convection at 20° and 30°C while data acquired at 10°C showed the complete absence of such signs of convection (data not shown).

Subsequent 10°C AAV SV-AUC runs using the Optima_4 as well as Optima_3 have all continued to show the absence of any data scan sharpening. This data strongly supports our initial conclusion that positive dT/dr gradients, generated via the Optima’s temperature control system, is the key factor for the observed convection disturbances seen in AAV SV-AUC runs made on the Optima. As a result of this data, conducting AAV SV-AUC experiments at 10°C (and if needed as low as 4°C) on an Optima offers a significantly pathway for eliminating or minimizing the impact of convection. In general, when using the Optima, such a general modification of experimental conditions should be consider when experiencing convection in SV-AUC experiments.

## Discussion

Recent developments in gene therapy over the last decade or so has drawn the renewed attention of many companies and researchers to the use of viral vectors (using various virus systems, e.g., adenovirus [44–48] and AAV [49–51]), along with completely synthetic particles, involving liposomes and nanoparticles [52–57], as drug (nucleic acid) delivery systems. Initially, interest in this area was predominately concerned with the delivery of DNA therapeutic material, however, interest has widened to include the use of RNA therapeutics [58] such as interference ribonucleic acids, RNAi [59,60], and messenger ribonucleic acids, mRNA [61,62] along with gene editing payload material [63–66]. Given the success and potential offered by these new classes of biopharmaceuticals, they are revolutionizing drug development. This has been clearly demonstrated today by the incredible rapid development of vaccines and therapeutic agents for dealing with the recent COVID-19 pandemic [67–70]. These and other events are driving the very near future appearance of a whole new array of more effective therapeutics. As a result, the landscape of biopharmaceutical development is progressing rapidly into areas where much larger and even more complex biopharmaceuticals are being produced [71]. In developing these very large and more complex drug product biopharmaceuticals scientists are being challenged to find adequate analytical tools to characterize them in terms of assessing their biophysical homogeneity (e.g., incompletely assembled or fragmented, empty, partial filled, completely filled, over filled and aggregated particles). Acquisition of such information is important in making sound and timely decisions in the development of these new biopharmaceuticals and eventually in determining their consistency of manufacturing by making sure that their biophysical attributes fall within established QC specifications for drug product release. Today the analytical capabilities of AUC can meet these challenges [17,18]. It is a golden opportunity for this instrument to shine and fill an important analytical void.

In this paper, work carried out on a number of different Optima units used in characterizing AAV drug delivery systems, a performance issue has been identified that is believed to be a remnant problem of the initial temperature issue uncovered when the Optima was first released into the field over 3+ years ago. This remaining temperature problem is believed to stems from the basic design of the temperature controlling system in the Optima, which was also very likely responsible for the introduction of a shroud within the rotor chamber (to disperse heat), along with some software changes associated with the temperature controller to avoid the initially encountered convection issue.

### Sources of heating and cooling during AUC

In considering temperature as the underlining source of the Optima’s present temperature-convection problem, other than the direct cooling and heating of the rotor via the Optima’s Peltier cells (in its temperature control system) the list of other sources of heating and cooling that can be encountered during AUC include the following:

1. The continuous transfer of heat from the rotor drive shaft to the AUC rotor as it is rotating [32–34, 36].
2. Cooling from the adiabatic physical stretching of the rotor during its acceleration to running speed [27,35,36].
3. Heating from direct radial adiabatic compression of the liquid sample in the AUC cell [36,72] and rotor compression around the outer bottom edge of the AUC cell during the acceleration of the rotor to running speed [36].
4. Frictional heating along the outer skin of the rotor due to the residual air still present inside the rotor chamber [32,33,36].

In the case of “1” the radial location of the drive yields a negative temperature gradient with respect to increasing radius. This leads to a positive density gradient that would actually provide stabilization against convection. In the case of “2” and “3”, it has been speculated that these two sources could work together to set up a positive temperature gradient with increasing radius to provide a negative (destabilizing) density gradient [36]. Such a situation could explain the convection seen in the Optima. However, the relatively low rotor speed and the low compressibility of the materials involved make this overall contribution to the temperature-convection problem very small. Finally, in the case of “4”, although the greatest frictional heating would likely take place at the highest radial region of the rotor where the linear velocity is the greatest, leading to positive a positive dT/dr, the relatively high vacuum used in AUC should greatly minimize this as a potential problem.

Although the above mentioned sources of heating and cooling, cases “2-4”, could contribute to the Optima’s temperature-convection problem, what is important to realize is that *all* these sources of heating and cooling also exist when using the XL series analytical ultracentrifuge. Consequently, if any of these sources of heating and cooling were the root source of the Optima’s temperature-convection problem, then using the same experimental running conditions in terms of temperature, rotor, rotor speed and rate of acceleration for conducting AAV SV-AUC experiments on the XL-I should also give rise to the same data scan sharpening artifact seen in the Optima. However, this is not the case. XL-I AAV SV-AUC experiments have not shown the presence of data scan sharpening. As a result, this rules out these other sources of heating and cooling as root cause of the Optima’s temperature-convection problem.

Consequently, the Optima’s temperature-convection problem revealed when studying AAV, appears to be rooted in the Optima’s temperature control system. Specifically, the underlining problem with this system is manifested by the physical (radial) location of the cooling/heating elements (Peltier cells) located along the outer wall of the rotor chamber. Such a physical arrangement leads to the generation of adverse positive radial temperature gradients, dT/dr gradients, especially when the Peltier cells are called upon to provide heat. Such temperature gradients will give rise to negative (destabilizing) dρ_T_/dr gradients with increasing radius that can propagate into the spinning sample solution. If these negative density gradients are sufficient such that the conditions indicated by eq. 6 in the Supplemental Material (for all appropriate r and t values during SV-AUC) cannot be met, convection will occur.

### Other critical factors to consider when dealing with the Optima’s temperature problem

It should be noted that the Optima’s convection problem discussed in this paper has surfaced under ultracentrifugation conditions that are particularly susceptible to convection. These conditions concern the analysis of dilute solutions of very HMW solutes where low rotor speeds are used [36, 73–76], as commonly employed in AAV SV-AUC work. In these cases a weakness in the strength of the radial density gradient (generated predominately by the common physiological, but relatively low levels of LMW components in the formulation buffer used in these studies) can exist (see the horizontal gray color bars in Figures 4A-D) enabling the presence of weak destabilizing perturbations to give rise to convection. However, given that data scan sharpening effects are not typically observed when Optima studies are conducted on other biopharmaceuticals (like mAbs) at much higher rotor speeds, where stronger formulation buffer density gradients are generated (see Figure 5), it would seem to indicate that the inherent temperature controlling problem on the Optima can be much better tolerated under higher rotor speeds. Consequently, running AAV at much higher rotor speeds would appear to be a relative simple solution for running AAV on the Optima to avoid the convection seen at much lower rotor speeds. However, when AAV SV-AUC experiments were simulated at a much high rotor speed, the results revealed that there was less density gradient protection from the formulation buffer than at lower rotor speeds (compare AAV vs formulation buffer data scan results in Figure 4 at 18k rpm, to the results for AAV vs formulation buffer data shown in Figure 2S at 40K rpm, Supplemental Material, *Increased potential for convection when conducting AAV SV-AUC at high rotor speeds*, noting that the Y-axis is the same in both).

Given the above situation what can be important to consider when choosing or planning to increase rotor speed, is the difference in sedimentation coefficient between the macromolecular solutes of interest and its formulation buffer components. When this difference in sedimentation coefficient, relatively speaking, is not that great (e.g., as in the case of mAbs vs formulation buffer components), enough time should be available for the formulation buffer components (typically present at physiological concentrations, totally about 0.1-0.2 M) to set up an adequate stabilizing density gradient (dρ/dr ≥ 0). At present these considerations can be particularly important when using the Optima, as shown for AAV work, given the configuration of the Optima’s present temperature control setup.

Thus given the above analysis, even at high rotor speeds the Optima’s temperature control system can still play a visible role in degrading data quality. Thus care needs to be given even when considering the use the Optima at high rotor speeds in trying to generate stronger formulation buffer density gradients to avoid convection, to improve resolution and when assessing aggregation of typical protein drugs (e.g., mAbs) when the aggregates that are presence are very high in MW (e.g., having MWs approaching that of the AAV monomer). In these cases the relative differential difference in sedimentation coefficient between the formulation buffer components and the macromolecular material of interest being studied can be sufficient to lead to situations where macromolecular material can migrate into a relative weak region of the stabilizing density gradient (generated by the formulation buffer material) that is similar to the situation shown in Figure 4, or worse, virtually absent, as seen in the case of Figure 2S shown in the Supplemental Material. In these situations, given the present state of the Optima’s temperature control system, convection may occur, leading to alterations in SV-AUC data.

### Possible experimental approaches for avoiding convection in the Optima

Approaches to potentially remedying the Optima problem, beyond that of making alterations to the instrument’s temperature control system, such as repositioning of the Peltier cells (heating/cooling elements of the Optima’s temperature control system), further changes in control software, improvement in the quality of key components or possibly further improvements in the heat dispersive properties of the shroud, consist of the following changes in experimental conditions:

1. Increase the concentration of the formulation buffer or just one of its LMW solutes or add an additional (appropriate) new LMW solute (e.g., such as sucrose [76,77]) to strengthen the stabilizing properties of the formulation buffer’s density gradient, dρ_B_/dr.
2. Intentionally induce a small negative temperature gradient across the rotor in order to generate a positive (stabilizing) density gradient. This might be done by choosing a starting temperature that is slightly higher than the final run temperature, then dropping the desired temperature down to the final run temperature shortly before acceleration. This ‘fix,’ however, may result in analysis problems.
3. Decrease the running temperature of the SV-AUV experiment to reduce the destabilizing negative density gradient generated by any temperature perturbation gradient, dρT/dT, by taking advantage of the reduced change in density of water as its temperature is reduced to achieve stable dρ/dr values (dρ/dr ≥ 0) to prevent convection (see Figures 7B and in Supplemental Material see Figure 1S, and discussion around eq. 6).

In case “1” the approach actually negates one of the strong reasons for using AUC, which is its ability to conduct characterization work on a sample in its actual *native* solution or formulation buffer that the sample is stored in. Such changes can potentially alter the physicochemical properties of the sample and thus provide misleading information, especially about aggregation. However, if this approach is pursued, ideally the LMW solute(s) chosen to increased should typically have the highest density increment, dρ/dc, to achieve the highest radial density gradient, see discussion in the Supplemental Material, *Key criteria for stable convection-free SV-AUC* concerning density increments.

In the case of “2”, one could actually take advantage of the physical location of the Peltier cells to generate a slight temperature gradient across the rotor by equilibrating the rotor at a temperature setting that is slightly higher than the desired running temperature, which on the initiation of the run is reduced to the actual desired running temperature. This approach is very much like the approach used on the Model E to control temperature and provide a slight stabilizing density gradient (which is also discussed in the Supplemental Material, *Detrimental effects of a positive radial temperature gradient during SV-AUC*). This approach, it should be noted, does give up a small uncertainty in the accuracy of experiments running temperature, but it may prove useful. At present this approach has not been investigated and is limited in the case of the Optima to only a maximum temperature offset of about 0.5°C.

In the case of “3” repetitive AAV SV-AUC runs on two Optima units, which demonstrated data scan sharpening at 20°C, have in fact shown the absence of any such data scan sharpening when run at 10°C. Given the common need to store AAV material at low temperatures, to achieve long term stability, it’s felt that the use of 10°C versus 20°C does not constitute a significant change in experimental condition that would draw concerns in potentially altering the characteristics of an AAV sample’s homogeneity. At present, this approach appears to be the simplest and most practical approach for dealing with the Optima’s temperature-convection problem.

### Final conclusion

The designing and construction of an analytical ultracentrifuge is not an easy task. The integration of multiple sub-systems into a complete well-functioning instrument is a challenge. In working with a number of Optima users an issue has been uncovered with one of these sub-systems, the temperature control system. In this paper key pieces of data are summarized that point to this sub-system’s problematic behavior as the source of the micro-convection problem that can give rise to data scan distrubances. In addition, this paper has provided the signs to look that reveal the presence of this problem and a temporary workaround for avoiding or minimizing the problem. Nevertheless, until this problem is permanently addressed, present and future users of the Optima need to be aware of this problem and to be able to recognize its presences within their data to enable them to better assess their data’s quality. Such information is of particular importance when employing the Optima in the biopharmaceutical industry, where important decision making needs to be made during drug development, and if used in the quality control (QC) environment as a QC analytical tool, for biopharmaceutical lot release.

## Acknowledgement

The authors would like to thank Dr. Peter Schuck for his helpful suggestions and comments on reviewing the manuscript.

## Supplemental Material

### Key criteria for stable convection-free SV-AUC

The movement of solute molecules in solution under the influence of a centrifugal field (during boundary SV-AUC) can provide significant information about the physical properties of these solute molecules. To acquire this information, however, requires that the migration of these solute molecules be studied in a convection-free environment (where transport of solute molecules does not occur due to bulk solution flow or movement). In order to successfully achieving this situation in an analytical ultracentrifuge the presence of an adequate radial density gradient, dρ/dr, needs to be in place with the following property, see eq. 1 below:

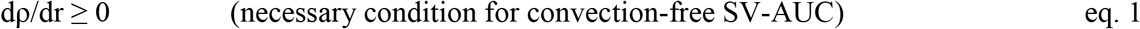

This above condition needs to be met for all radial positions, r, along the entire sample solution column, from the sample sector solution meniscus (at the air/liquid interface) to the sample sector cell’s base, over the entire time duration, t, of the SV-AUC experiment, from the start of the rotor’s acceleration to the end of useful data acquisition.

Although successful SV-AUC, as noted in eq. 1, can be achieved under conditions where dρ/dr = 0, such a condition is meta-stable. Under this particular condition migration of solute molecules in a gravitational field is in a very precarious state. Subtle (local) adverse perturbations in solution density (e.g., due to local changes in temperature or/and sample concentration) can result in convection. Consequently, it is far preferable to have dρ/dr > 0 where the greater the magnitude of dρ/dr the less is the opportunity for encountering convection. Nevertheless, dρ/dr should not be that great that its variation with r and t itself significantly influences the migration of sample solute molecules during the sedimentation process. If this latter situation exists the resulting analysis will be more complex requiring the variation of density and viscosity with respect to both r and t to be accounted for when trying to appropriately model the sedimentation process [1].

In conducting boundary SV-AUC Schachman [2] pointed out there are two general classes of components that contribute to dρ/dr. These two components include the radial density gradient formed from sample macromolecular solutes (S) being studied, dρ_S_/dr, and the collection of low molecular weight, LMW, solutes or excipient molecules (salts, buffer, sugars, etc.) in sample solution that constitute the formulation buffer (B) of the sample solution, dρ_B_/dr. Thus, based on eq. 1 the condition for convection-free, now becomes eq. 2 below:

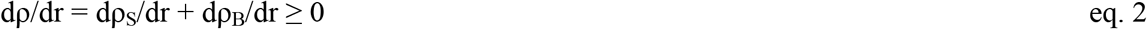

In the case of the formulation buffer each solute contributes its own radial density gradient contribution to the total radial density gradient, which is equal to the change in the component’s concentration with respect to radius (dc/dr), multiplied by its respected density increment (dρ/dc, which is equal to (1 - Φ’ ρ_o_) [3], where Φ’ = apparent partial specific volume of the solute and ρ_o_ = density of the solution the solute is dissolved in), which for all practical purposes is a constant for any solute over a wide concentration range. The density increment for each solute is a unique physicochemical property of that solute in solution, where the larger its value the greater the density gradient that the solute can form per unit concentration in the same gravitational field (note, since formulation buffers typically are composed of several solutes, increasing the concentration of the solute with the highest density increment would be an optimal approach for achieving the highest dρ_B_/dr value to insure convection-free AUC, e.g., if a formulation buffer contains Pluronic, dρ/dc ≈ 0.1 [4], and sucrose, dρ/dc ≈ 0.4 [5], increasing sucrose over Pluronic would be optimal assuming there are no stability issues associated with macromolecular material). Hence eq. 2 becomes eq. 3:

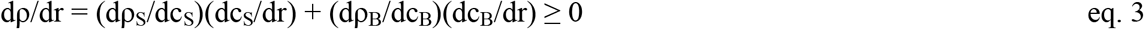

In the case of AAV work, however, AUC analysis is typically carried out under very dilute AAV concentration, leading to very low value for dcS/dr. As a result its contribution to the overall density gradient, dρ/dr, is negligible (dρ_S_/dr ≈ 0) compared to dρ_B_/dr. Consequently for AAV SV-AUC experiments the key stabilizing radial density gradient is effectively dependent on the radial density gradient provided by the formulation buffer, dρ_B_/dr, hence eq. 3 simplifies to eq. 4:

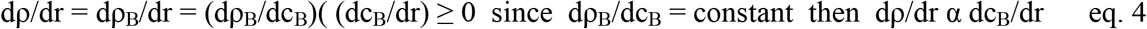

In conducting SV-AUC one might also think that the resulting pressure gradient, formed during AUC due to its inherent variation with radius within a centrifugal field (resulting in a corresponding variability in sample solution compressibility) would also be a direct contributing factor to dρ/dr. However, Pickel in 1942 [6] showed this was not the case. Nevertheless, solution compression can induce an indirect impact on solution density through the release of heat (and thus change in temperature) during the rotor’s acceleration to running speed [7]. Indeed, the indirect effect of temperature on solution density (as well as viscosity) can play a critical role in achieving or not achieving accurate and stable SV-AUC. As a result, SV-AUC needs to be conducted under constant temperature (to avoid the potential negative impact of temperature changes on the density and viscosity of a sample solution during AUC). Details on how the Optima goes about controlling temperature play a critical role in this paper in exposing samples run on the Optima to convection.

### Detrimental effects of a positive radial temperature gradient during SV-AUC

In assessing the impact of temperature gradients on the sedimentation of solute molecules in an aqueous solution during SV-AUC, one needs to realize that as temperature increases the density of an aqueous solution decreases, see Figure 1S for the case of distilled water. Conversely, as the temperature of an aqueous solution is decreased its density increases until the temperature approaches a value of about 4°C, where the unique properties of water make this no longer true, see Figure 1S.

**Figure 1S.**
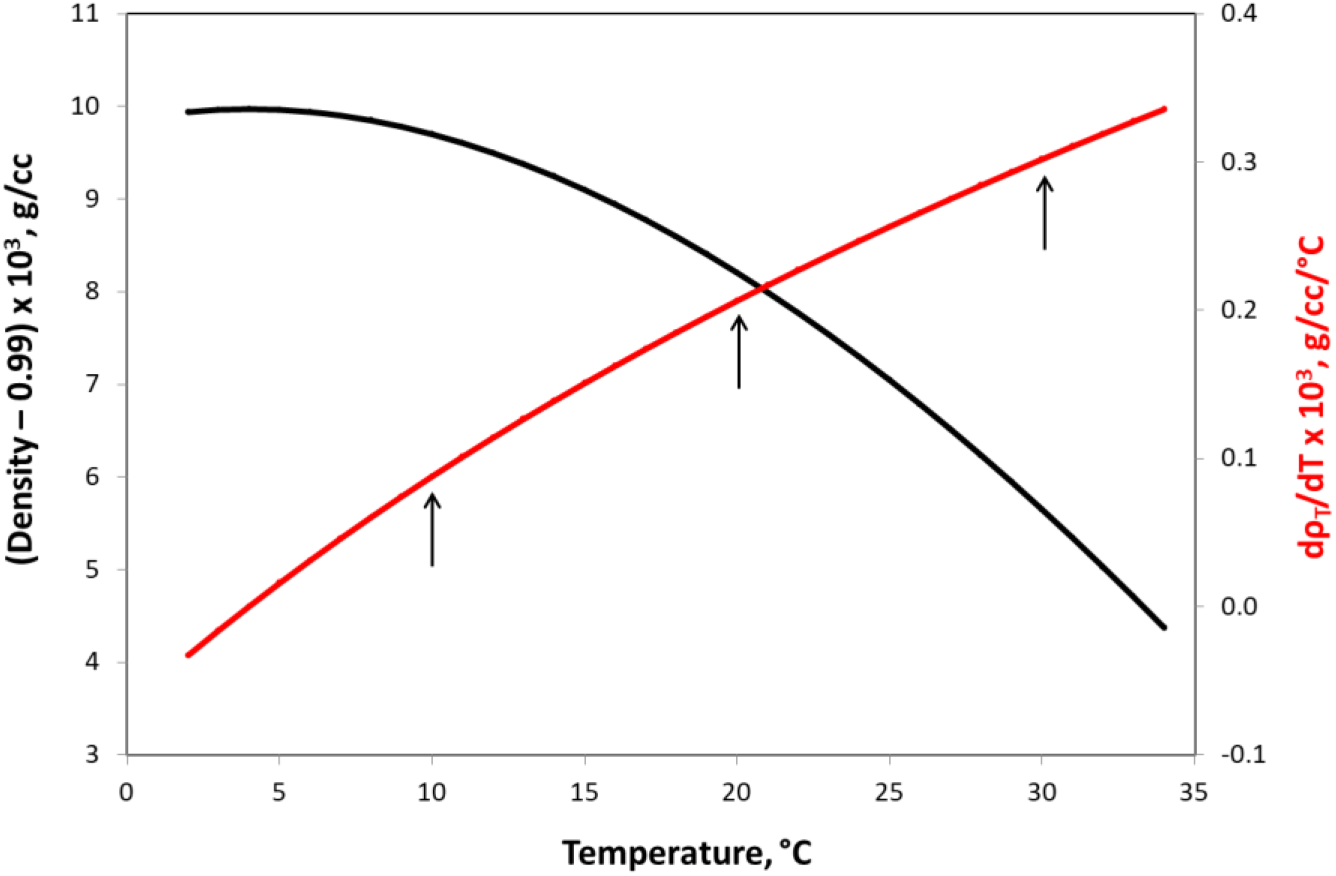
A plot of the density of distilled water as a function of temperature [8] and its associated plot of dρ/dT as a function of temperature. dρ/dT values were approximated by computing Δρ/ΔT values from the density versus temperature data, which are spaced every 1 degree, by evaluating the change in density (Δρ) between two adjacent temperatures (where ΔT is a constant equal to 1). Each computed dρ/dT ≈ Δρ/ΔT = Δρ value was then assigned to the higher of the two adjacent temperature values used in assessing Δρ. Vertical arrows shown correspond to 10, 20 and 30°C.

As is indicated in eq. 1 convection-free sedimentation in an analytical ultracentrifuge requires a radial solution density gradient, dρ/dr, that is ≥ 0 (preferably > 0) to be present in the sample solution held within the sample sector of the AUC cell over all appropriate r and t values of a SV-AUC experiment. The presence of a positive temperature gradient with respect to increasing radius will contribute a negative (destabilizing) density gradient, dρ_T_/dr, with respect to increasing radius. If this situation is allowed to be present during SV-AUC it can cause convection, due a density inversion as indicated in Figure 1 in the main body of the paper, unless a greater opposing (positive) radial density gradient with increasing radius can be generate to override it. As a result positive radial temperature gradients must be avoided during SV-AUC, see eq. 5a. On the other hand the presence of a negative temperature gradient with respect to increasing radius will contribute a positive (stabilizing) dρ_T_/dr with respect to radius providing a protective density gradient, which can help achieve convection-free SV-AUC, see eq. 5b:

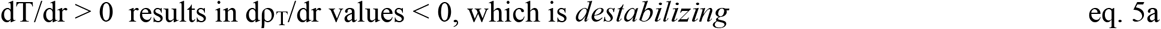

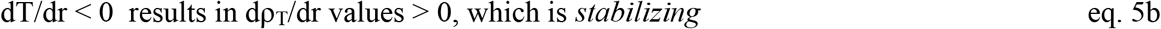

As a result, when conducting SV-AUC on dilute samples, where the only significant stabilizing density gradient is from the LMW components in the sample’s formulation buffer (dρ_B_/dr, see eq. 4), if a positive temperature gradient does get introduced into a dilute sample during SV-AUC (as shown in Figure 7B in the main body of the paper) in order to achieve stable convection-free sedimentation eq. 6 must be true (for all appropriate r and t values):

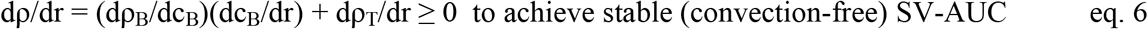

Consequently, eliminating or reducing negative dρ_T_/dr values so the criteria indicated by eq. 6 can be achieved is critical.

By looking more closely at dρ_T_/dr we can show an interesting opportunity for minimizing the impact of temperature on this problem, by realizing dρ_T_/dr is actually composed of two factors, see eq. 7:

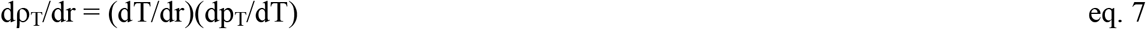

The first term, dT/dr, is the temperature gradient (that may exist in a sample during the course of SV-AUC), which as discussed above, ideally needs to be equal or less than zero. Attempts at achieving this goal is predominately the role of a well-designed functioning temperature control system in an analytical ultracentrifuge. The second term, dp_T_/dT, is a factor that is a physiochemical property of the test sample’s matrix, which in the case of AAV SV-AUC work is an aqueous solution, offering a very interesting opportunity for minimizing the impact temperature on dρ/dr. It turns out in the case of water dρ/dT (which should effectively be equivalent to dp_T_/dT) actually decreases with temperature, reaching a value of zero at about 4°C, see Figure 1S. Consequently, by lowering the running temperature of a SV-AUC experiment one can significantly reduce (or eliminate) dρ_T_/dr contribution to eq. 6. As a result, simply reducing the running temperature of a SV-AUC experiment can significantly take the burden off both the analytical centrifuge’s temperature control system and the formulation buffer’s density gradient to successfully achieve convection-free SV-AUC.

### Increased potential for convection when conducting AAV SV-AUC at high rotor speeds

It might seem reasonable to expect that a higher rotor speed would result in a greater gradient, thus preventing convection. However, simulation of a AAV SV-AUC run at 40K rpm at 20°C, Figure 2S shows that the increased speed actually may *increase* the opportunity for convection due to the wider radial window where the density gradient is weak (e.g. compare this simulation to that for 18K at 20°C, Figure 4 in the main body of the paper, noting also the weaker density gradient provided by the formulation buffer in this radial window in the case of 40K vs 18K).

**Figure 2S.**
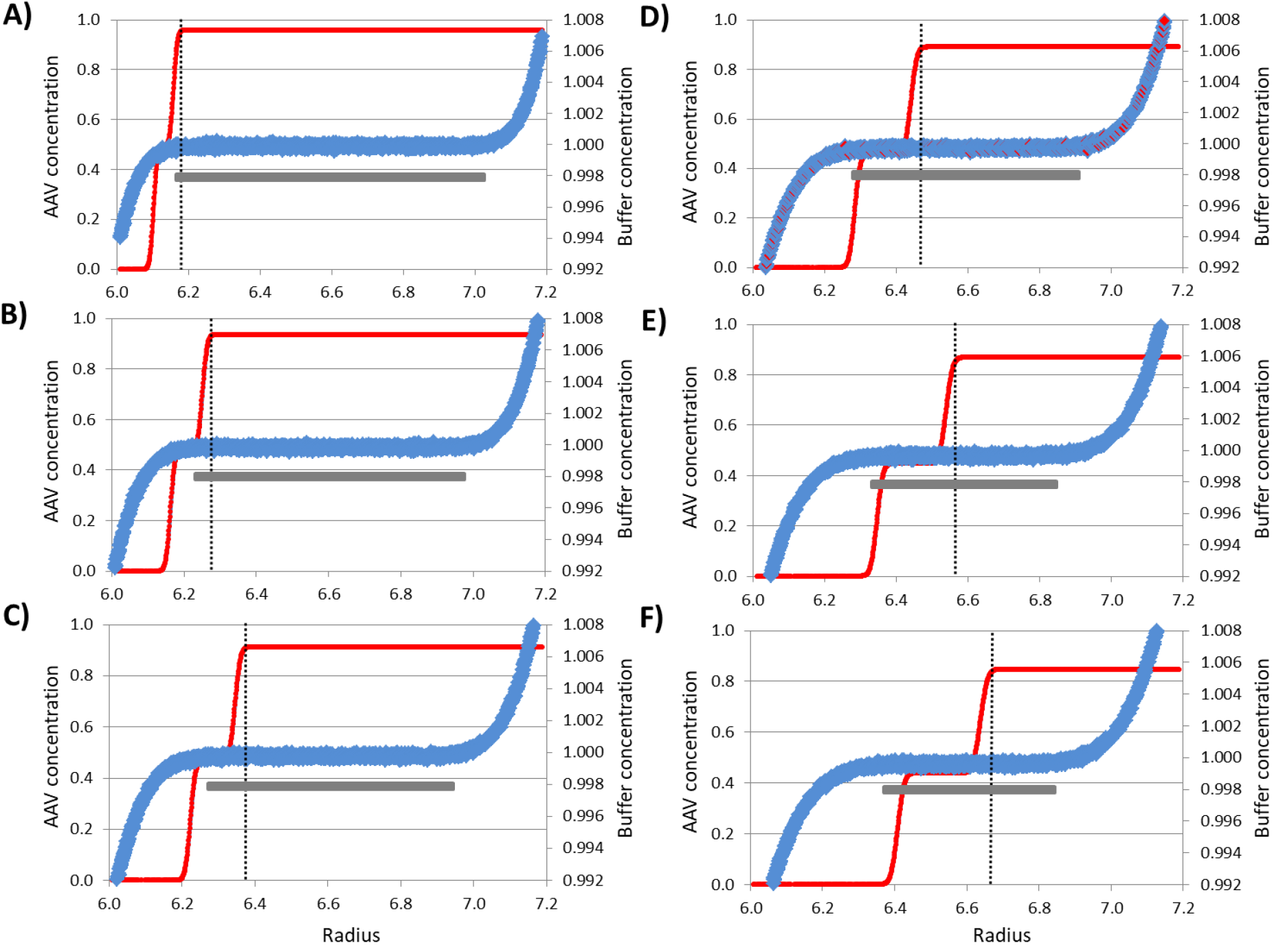
Simulated AAV SV-AUC data for a hypothetical AAV sample consisting of 50:50 a mixture of empty and full virus particles and for its formulation buffer, as outlined in Figure 4 in the main body of the paper, but at 40K rpm instead of 18K rpm. Data scans for the AAV mixture and formulation buffer were calculated for times intervals of 90 secs resulting in data scans for the following time points (from the start of the run): **A)** 90 sec, **B)** 180 sec, **C**) 270 sec, **D**) 360 sec, **E)** 450 sec and **F)** 540 sec for the same radial positions. Black vertical dotted line corresponds to the position of the leading boundary edge of the full AAV material. In both cases the initial starting concentration was an arbitrary concentration of 1. The length of the horizontal grey bars included in these plots indicates the extent of the radial region where the density gradient from the formulation buffer is the shallowest (where the potential for convection is most significant); while the lighter the grey the less prominent is the instability (or weakness) in this density gradient. In this case note that the level of instability and the reduction in radial region where potential instability exists are both only moderately reduced over the time course of this SV-AUC simulation.

